# Neurogenesis mediated plasticity is associated with reduced neuronal activity in CA1 during context fear memory retrieval

**DOI:** 10.1101/2021.03.24.436893

**Authors:** Alexandria Evans, Dylan J. Terstege, Gavin A. Scott, Mio Tsutsui, Jonathan R. Epp

## Abstract

Adult hippocampal neurogenesis has been demonstrated to affect learning and memory in numerous ways. Several studies have now demonstrated that increased neurogenesis can induce forgetting of memories acquired prior to the manipulation of neurogenesis and, as a result of this forgetting can also facilitate new learning. However, the mechanisms mediating neurogenesis-induced forgetting are not well understood. Here, we used a subregion-based analysis of the immediate early gene c-Fos as well as in vivo fiber photometry to determine changes in activity corresponding with neurogenesis induced forgetting. We found that increasing neurogenesis led to reduced CA1 activity during context memory retrieval. We also demonstrate here that perineuronal net expression in areas CA1 is bidirectionally altered by the levels or activity of adult generated neurons in the dentate gyrus. These results suggest that neurogenesis may induce forgetting by disrupting perineuronal nets in CA1 which may otherwise protect memories from degradation.

## Introduction

The proliferation and integration of new neurons in the adult mammalian brain is a unique form of plasticity that has the potential to both create new connectivity as well as to disrupt existing circuits. As a consequence, there are numerous implications for adult neurogenesis as a modulator of hippocampal structure and function. Previous work has demonstrated that, in many cases, elevated levels of neurogenesis positively impact processes such as learning^1–3^, memory^4,5^ and, cognitive flexibility^3,6^. Recent work has also shown that in addition to the positive influences that neurogenesis may have on learning and memory, there is also a weakening of memories that were formed prior to the elevation of neurogenesis^7–14^. Ultimately, this role of adult neurogenesis is also beneficial as it mitigates proactive interference between old and new memories. This occurs particularly in situations where there is conflict between the two memories, such as in the case of reversal learning. As a result, neurogenesis induced forgetting is beneficial for learning as it allows for faster acquisition of new memories^9^. This “retrograde” effect of neurogenesis impacts a wide variety of memory types, occurs in both females and males and, occurs regardless of how neurogenesis levels are increased (i.e. genetic, chemical, physical methods). Conversely, decreased neurogenesis reduces forgetting of previously acquired memories and slows the acquisition of new conflicting memories^9^. Together, these results demonstrate the importance of adult neurogenesis for maintaining the balance of stability versus plasticity and how this may preferentially modulate learning versus memory.

Neurogenesis induced forgetting is a robust and reliable phenomenon. However, the mechanisms that mediate the reduction in memory strength have not been clearly identified. New neurons integrate into the existing hippocampal circuitry and as result may overwrite existing mature synapses^15,16^. As such, it has been proposed that neurogenesis causes forgetting via reconfiguration of the mossy fiber to CA3 circuitry, as newly born DG cells form connections, leading to a failure to reactivate the original memory trace^7^. Computational modelling of the effects of neurogenesis on hippocampal circuitry has found that the addition of new, more excitable units to the DG after the network has been trained on a given pattern causes a reduction in the rate of reactivation of CA3 units that were previously activated by that pattern^17^. Indeed, ablation of neurogenesis impairs the reactivation of engram cells in the CA3 during fear recall^18^, showing that newly born DG cells have a significant effect on the activity of the CA3 during memory retrieval.

Computational models and predictions about potential mechanisms have, to date, been heavily focused on the DG and its important interactions with immediate input and output structures (e.g. entorhinal cortex and CA3^19–21^). Early in their maturation, adult-born granule cells preferentially synapse onto CA3 interneurons rather than excitatory neurons^22^ meaning that the addition of new neurons may not only have the effect of “overwriting” older mossy fiber connections, but also of causing a broad increase in inhibitory drive within the CA3. Although effects in the CA3 almost certainly play a role, neurogenesis has been shown to have downstream effects beyond the mossy fiber pathway. For example, increased neurogenesis causes increased inhibition in the CA3 and CA1^15,20^. Given that the Schaffer collateral pathway has denser connectivity than the mossy fiber pathway^23–26^, neurogenesis-induced changes in CA3 activity could lead to even larger changes in CA1 activity. Thus, we sought here to examine the impact of adult neurogenesis on activity of the DG, CA3 and CA1.

We hypothesized that adult neurogenesis impairs the retrieval of previously acquired memories by altering the activity or excitability of downstream hippocampal subregions. We tested this hypothesis by examining neurogenesis-dependent changes in the activity of hippocampal subregions during memory retrieval. Our results point to altered CA1 activity that may be mediated by the deterioration of perineuronal nets as a mechanism underlying neurogenesis induced forgetting. These findings provide a mechanistic account of the retrograde effects of adult neurogenesis and also help to reconcile the pro-mnemonic anterograde effects of adult neurogenesis with the pro-forgetting effects observed in the retrograde direction.

## Results

### Voluntary Running Increases Neurogenesis and Decreases Contextual Fear Memory Retrieval

Our first goal was to confirm the impact of voluntary exercise on memory retention in the retrograde direction (Fig. 1a). To do so, we first trained mice to establish contextual fear memory (Fig. 1b). To increase neurogenesis, half of the mice were given a running wheel for 4 weeks and the other half remained sedentary in standard housing conditions. Following this manipulation, we observed a significant decrease in memory retrieval in the running group (Fig. 1c) consistent with previous reports that elevated neurogenesis induces forgetting. To confirm that running increased neurogenesis, we examined the immature neuron marker DCX and observed a significant increase in the number of labeled cells in the dentate gyrus of runners (Fig. 1d,e). To determine whether the disruption of contextual memory following voluntary exercise is specific to memories acquired prior to the manipulation, we performed an anterograde style experiment where mice were given access to a running wheel prior to contextual fear conditioning (Supplemental Fig. S2a). We then tested the mice 30 days later. Despite the prior period of voluntary exercise, we observed no significant differences between control and running groups in contextual fear memory acquisition (Supplemental Fig. S2b) or subsequent retrieval (Supplemental Fig. S2c). Together these results demonstrate that voluntary exercise induces a large increase in neurogenesis and strongly modulates the strength of existing hippocampus-dependent memories.

**Figure 1.**
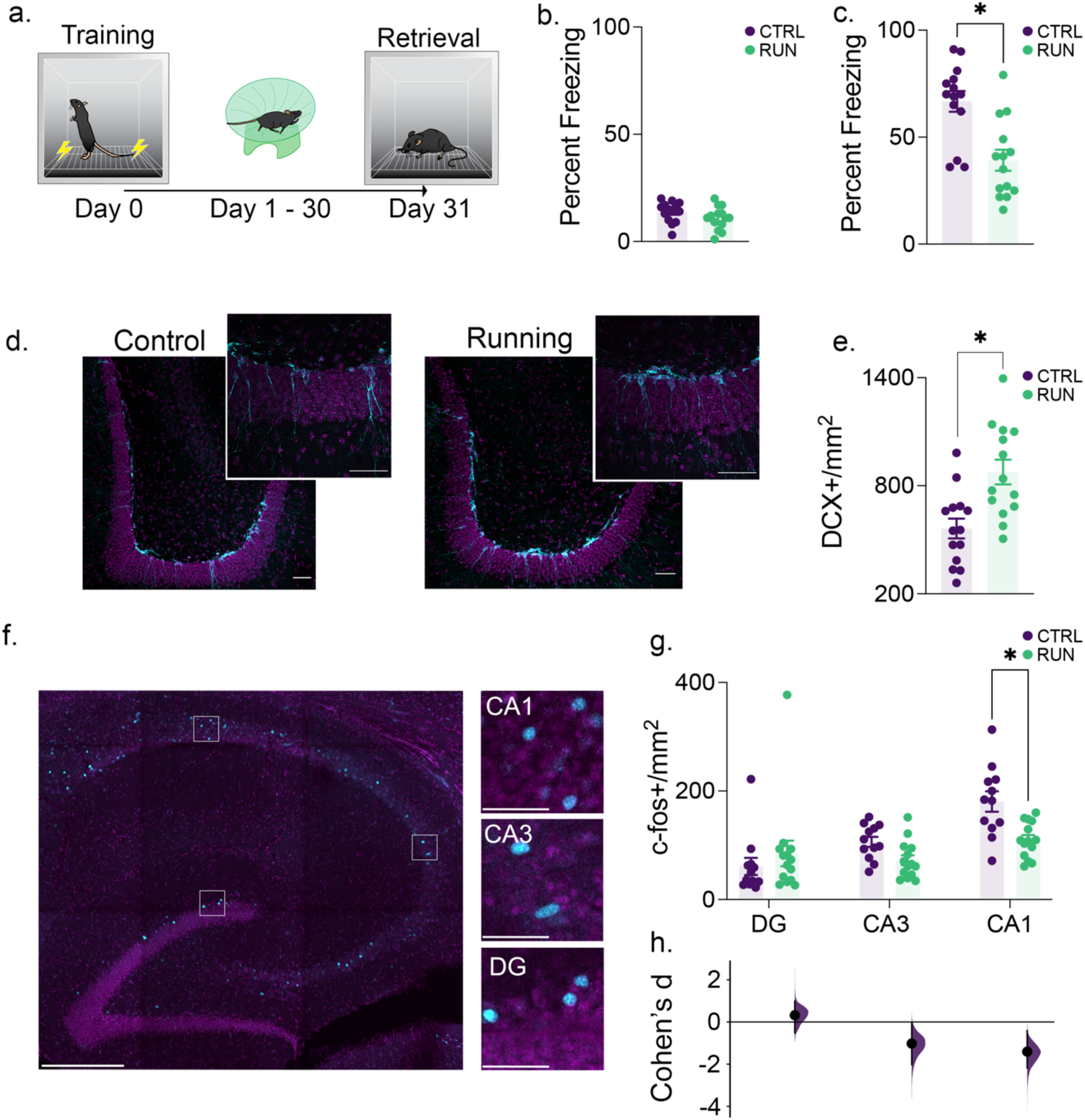
Running induces neurogenesis, promotes forgetting of previously acquired information and alters CA1 activity. (**a**) After training in a contextual conditioning paradigm, mice were housed conventionally (*n* = 14) or given running wheel access (*n* = 14) for 30 days before testing memory retention. (**b**) There was no difference in the proportion of time spent freezing between groups during training. (**c**) Running promoted forgetting of the conditioned context as measured by a decrease in freezing compared to sedentary control mice. (**d**) Example of doublecortin labeling (cyan) in the dentate gyrus of mice from control and running groups. Scale bars, 50 μm.(**e**) Mice given access to running wheels showed an increase in neurogenesis. (**f**) Examples of c-fos+ neurons (cyan) in hippocampus subregions during contextual fear conditioning memory retrieval. Hippocampus overview scale bar, 500 μm. High magnification scale bar, 50 μm. (**g**) Running did not alter the density of c-fos+ neurons in the DG or CA3 but increased c-fos expression in area CA1. (**h**) This alteration is further illustrated in a plot of the effect size (Cohen’s d), showing that there was only a condition-dependent decrease exceeding the bootstrapped 95% CI in CA1. Data analysis used Two-Sample T-Test (**b**,**c**,**e**), ANOVA (**e**) with Tukey’s *post-hoc* test, and Multiple Two-Groups estimation statistics with Cohen’s d as a measure of effect size (**h**). \**P* < 0.05. Data shown are mean ± s.e.m. See Supplemental Table 1 for Full statistical analysis.

### Memory induced c-Fos expression is decreased in area CA1 following voluntary exercise

Several previous studies^7–14^ have demonstrated that the forgetting of previously acquired memories is induced by elevating the rate of adult neurogenesis and our current results confirm these previous findings. Our next goal was to determine whether we could observe disrupted patterns of hippocampal activity that underlie the neurogenesis induced forgetting as a first step towards determining the mechanism by which memories are weakened. To do so, we examined c-fos expression, as a proxy of activated neurons^27–31^, in different hippocampal subregions. We observed a significant group by region interaction in c-fos expression (Fig 1 f-g). There were no significant differences in the density of c-fos expression in the dentate gyus or area CA3. However, the running group had a significant decrease in c-Fos in area CA1 compared to the control group.

While the difference was significant only in CA1, we did observe a trend towards a decrease in c-fos expression in CA3 as well. To further analyze these changes, we performed estimation statistics to examine the effect sizes in each subregion. Interestingly, there appears to be a gradient of effect that increases from the DG, to CA3 and CA1 (Fig. 1f-g) suggesting that the impact of adult neurogenesis is smaller at the site of its physical integration than it is downstream in the hippocampal circuitry.

We also examined c-Fos expression from mice in anterograde condition where neurogenesis was modulated by voluntary exercise prior to memory acquisition. As discussed above there was no memory impairment in the anterograde condition as a result of running. Additionally, in this condition there was no effect of running on c-Fos expression in any of the hippocampal subregions during subsequent memory retrieval (Supplemental Fig. S2d,e). This strongly supports the notion that the reduction in CA1 c-Fos in the retrograde condition is a consequence of the decreased memory retention.

### GCaMP7f photometry in CA1 is attenuated during memory retrieval by voluntary exercise

Having identified an activity dependent signature of impaired memory retrieval in area CA1, we next performed fiber photometry in this region to develop a more nuanced understanding of the activity changes related to neurogenesis induced forgetting. The same behavioural paradigm and timeline was used as above except that we virally expressed GCaMP7f^32^ to monitor the population activity of area CA1 during contextual fear learning and post running memory retrieval (Fig. 2a-b). As shown in our c-Fos experiments (Fig. 1), we demonstrate first that voluntary running increased neurogenesis (Fig. 2c-d) and, to the extent that neurogenesis was increased by voluntary running, we observed a significant decrease in freezing (Fig. 2e). We also found a significant correlation between neurogenesis and memory retention in runners (Fig. 2f). Prior to the voluntary exercise manipulation, we did not observe any group differences in contextual fear acquisition or in any fiber photometry related metrics during training (Supplemental Figure S4). Following one month of running, we performed photometry recordings in a clean mouse cage to determine whether running had a non-specific (i.e., non mnemonic) effect on CA1 population activity. There was no significant difference between runner and sedentary groups with respect to baseline calcium activity (Fig. 2g) indicating that voluntary exercise does not cause a general shift in CA1 activity. We then immediately transferred the mice to the contextual fear chambers. Upon doing so we observed an increase in GCaMP7f activity in the control group that was significantly attenuated in the runners (Fig. 2h). We calculated the area under the curve for each photometry trace and observed that this metric was significantly correlated with memory strength as measured by time spent freezing (Fig. 2i,j). These initial photometry results are indicative of reduced population activity in CA1 following voluntary exercise that is specific to memory retrieval. This is consistent with the c-Fos expression changes shown in Figure 1.

**Figure 2.**
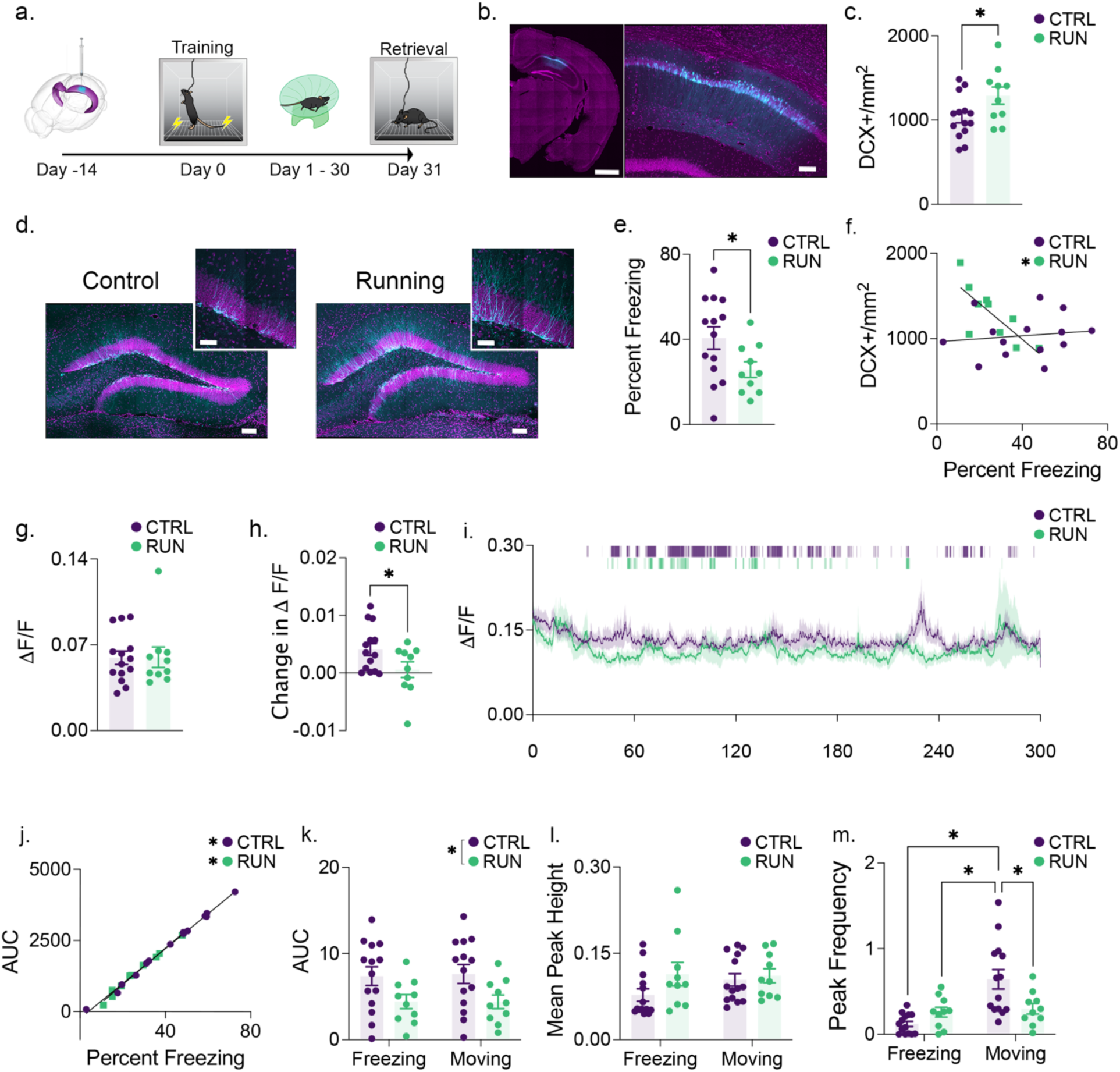
Running-induced neurogenesis increases behaviour-specific activity of CA1. (**a**) Mice were infected with GCaMP7f prior to contextual fear conditioning. Mice were then housed conventionally (*n* = 14) or with a running wheel (*n* = 10) for 30 days prior to a memory test. Fiber Photometry was performed during training and memory retrieval. (**b**) Representative photomicrograph of GCaMP7f expression (cyan). Overview scale bar, XX um. CA1 scale bar, 100 μm. (**c**) Runners showed an increase in neurogenesis. (**d**) Example of doublecortin labeling (cyan) in control and runners. Dentate overview scale bar, 100 μm. High magnification scale bar, 50 μm. (**e**) Running promoted forgetting of the context memory. (**f**) In runners, the increase in doublecortin labeling correlated with memory strength. (**g**) During photometry recordings in a clean home cage prior to the retention test, there was no difference in mean GCaMP7f activity between runners and controls. (**h**) Once transferred to the conditioning chamber, control mice showed a significantly greater increase in mean GCaMP7f activity from one context to the next compared to runners. (**i**) Mean GCaMP7f fluorescence across the context memory test with the median group freezing plotted above. See Supplemental Figure S3 for non-baseline-corrected group mean traces and representative individual recordings. (**j**) In both groups the area under the GCaMP7f fluorescence curve across the trial correlated significantly with percent freezing. Photometry recordings were then separated by behavioural expression. (**k**) Area under the curve was suppressed in runners irrespective of behavioural expression. (**l**) The mean peak height of the photometry signal did not differ across conditions or by behavioural expression. (**m**) Control mice showed an increase in GCaMP7f peak frequency during movement relative to freezing. Runners failed to show this behaviour-specific increase in frequency during movement. Data analysis used Two-Sample T-Test (**c**,**e**,**g**,**h**) and Two-way ANOVA (**k**,**l**,**m**) with Tukey’s *post-hoc* test. \**P* < 0.05. Data shown are mean ± s.e.m. See Suplemental Table 2 for full statistical analysis.

We were next interested in determining whether we could differentiate control versus running group based on differential CA1 activity during freezing versus non-freezing behaviours. We examined AUC and observed a significant decrease in runners (Fig. 2k). However, this was consistently decreased during both movement and freezing. We also analyzed the Peak frequency and while there was no significant main effect of group, there was a significant group by behaviour interaction. We examined the mean peak height and found that this metric was not significantly different between groups and also did not change as a function of freezing versus non-freezing behaviour (Fig. 2l). During bouts of freezing, both controls and runners showed low peak frequency. During bouts of movement, control mice exhibited a significant increase in CA1 peak frequency that was not observed in the running group (Fig. 2m). This suggests that the behaviour dependent differences in peak frequency are most likely not due to the recruitment of additional neurons (in which case an increase in peak height would be expected) but instead due to changes in the frequency of the same population of neurons^33^.

### Voluntary exercise modifies perineuronal net expression in the hippocampus

Perineuronal nets (PNNs) have proposed roles in modulating plasticity, excitability and memory strength.^34–36^ Therefore, we speculated that one potential factor that might be altered by enhanced adult neurogenesis and could potentially explain the associated decrease in memory expression could be the expression pattern of PNNs. We therefore quantified the density of PNNs in the DG, CA3 and CA1 of mice that had undergone post-learning manipulation of neurogenesis with voluntary exercise (Fig. 3a). Intriguingly, we observed the same pattern of changes as was observed with c-Fos expression, that is, a running by region interaction with a reduction of PNN density in the CA1 of the running group (Fig. 3b, c). Using high magnification confocal microscopy, we examined the surface area of the remaining PNNs in the CA1 and found that, in addition to the decrease in overall number, there was also a decrease in the contiguity of the surface structure in the running group, potentially indicative of degradation of the PNNs (Fig. 3d-f). Interestingly, to examine the extent that perineuronal net expression is disrupted by neurogenesis *per se*, we correlated both the contiguity and density of PNNs with the number of DCX-labeled immature neurons. In both cases, there were strong inverse correlations in the control group (Significant in the case of contiguity and a near significant trend with PNN density) whereas these correlations were weak and non-significant in the running group (Fig. 3g,h).

**Figure 3.**
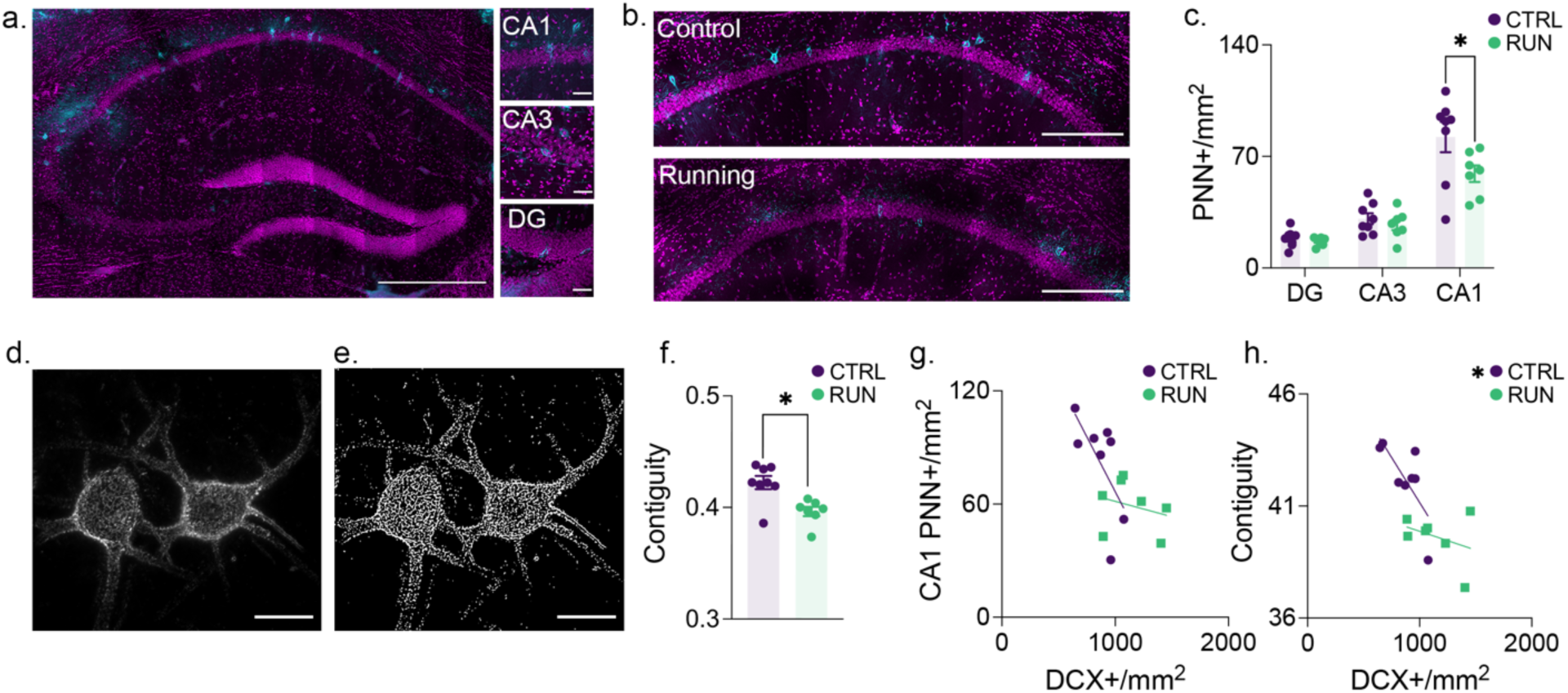
Running-induced neurogenesis decreases the density and contiguity of perineuronal nets in CA1. (**a**) Representative photomicrograph of PNN expression (cyan) across the hippocampus, with high magnification images of CA1, CA3, and the dentate gyrus. Hippocampus overview scale bar, 500 μm. High magnification scale bar, 50 μm. (**b**) PNN expression density in controls (*n* = 8) and runners (*n* = 7). Scale bars, 250 μm. (**c**) Running-induced neurogenesis decreased the expression density of PNNs in CA1. (**d**) High-resolution images of PNNs were collected from the CA1 and (**e**) binarized using thresholding to assess contiguity. Scale bars, 25 μm. (**f**) Running-induced neurogenesis decreased the contiguity of PNNs in CA1. In the control condition, the density of doublecortin+ cells trended highly in the anti-correlated direction with (**g**) CA1 PNN expression density and (**h**) CA1 PNN contiguity, while neither of these correlations were significant in the running condition. Data analysis used ANOVA (**c**) with Tukey’s test during *post-hoc* multiple comparisons and Two-Sample T-Test (**f**). \**P* < 0.05. Data shown are mean ± s.e.m. See Supplemental Table 3 for full statistical analysis.

### Adult Neurogenesis Modulates Perineuronal Net expression in CA1

To further determine whether neurogenesis is indeed the factor that is modulating the expression of perineuronal nets in CA1, we investigated additional regulators of adult neurogenesis. As an additional mechanism of increasing neurogenesis, we treated mice with memantine^10^ and to decrease neurogenesis we used temozolomide^37^ (TMZ). Compared to saline injected mice, memantine treatment resulted in a significant decrease in PNN expression in CA1 (but not DG or CA3), just as was observed with voluntary running. TMZ, on the other hand, reduced neurogenesis and subsequently increase the expression of PNNs in CA1 (Fig. 4a,b).

**Figure 4.**
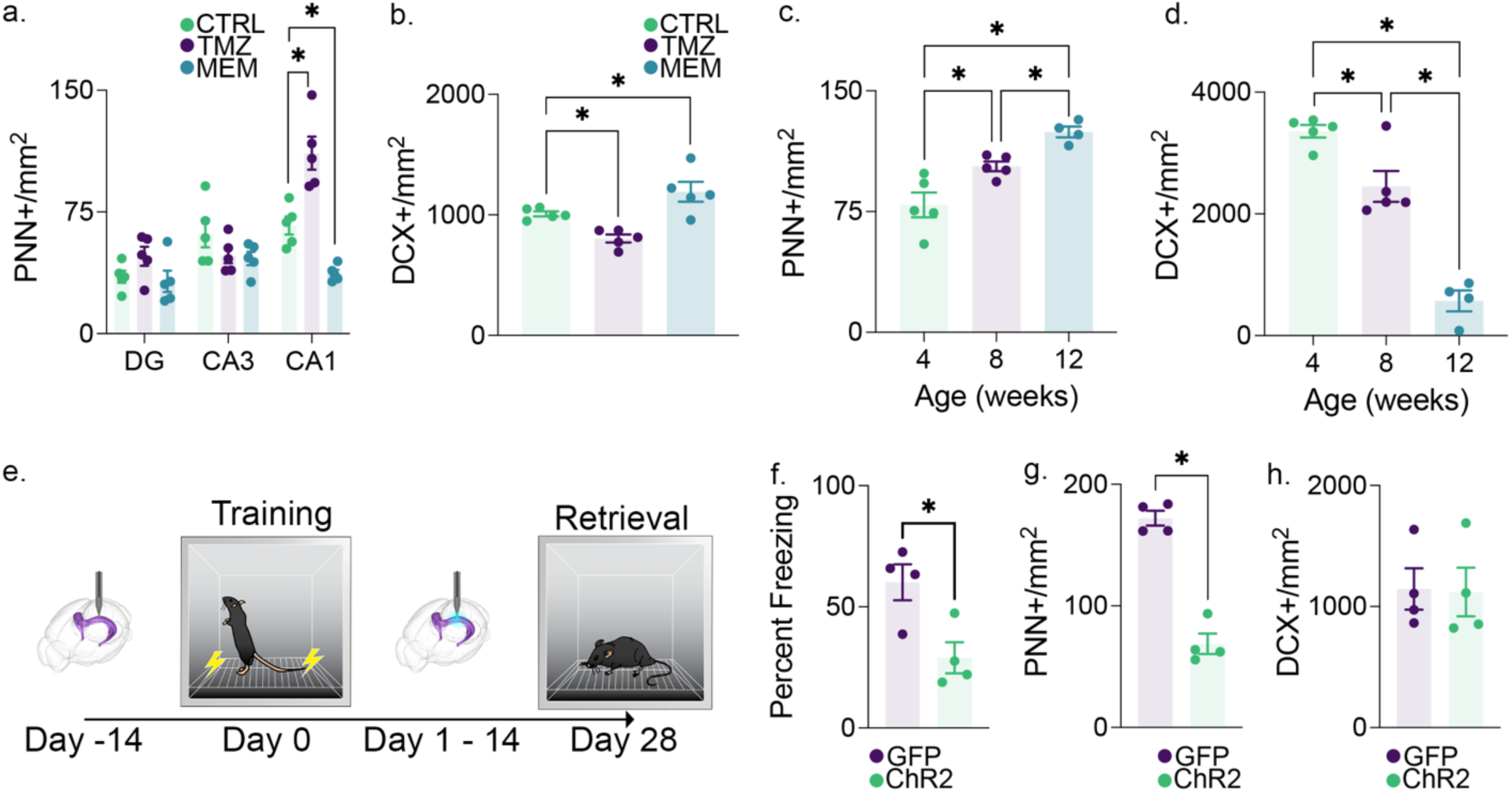
The activity of immature granule cells in the dentate gyrus is inversely related to the density of perineuronal nets in CA1. (**a**) 4 weeks of TMZ treatment (*n* = 5) increased the expression density and memantine treatment (*n* = 5) decreased the expression density of PNNs in CA1 relative to saline-treated controls (*n* = 5). This treatment effect was not present in the dentate gyrus or CA3. (**b**) Memantine treatment induced an increase in neurogenesis, while TMZ treatment reduced neurogenesis relative to saline-treated controls. (**c**) The expression density of PNNs in CA1 increased with age throughout adolescence, during which time (**d**) neurogenesis decreased. (**e**) Optic fibers were implanted in the dentate gyrus of tg+ (*n* = 4) and tg-(*n* = 4) Nestin-ChR2 mice 14 days prior to contextual conditioning. Nestin-expressing immature neurons in the dentate gyrus were then stimulated for 5 minutes daily at 10 Hz for 14 days. Mice were reintroduced into the conditioned context 28 days after conditioning.(**f**)During the reintroduction to the conditioned context, ChR2 tg+ mice showed impaired memory relative to ChR2 tg-controls. (**g**) ChR2 tg+ mice also showed a significant decrease in CA1 PNN expression density. (**h**) These changes in memory retrieval and CA1 PNN expression density occurred in the absence of differences in the expression density of doublecortin+ immature neurons. Data analysis used ANOVA (**a**,**b**,**c**,**d**) with Tukey’s test during *post-hoc* multiple comparisons and Two-Sample T-Test (**f**,**g**,**h**). \**P* < 0.05. Data shown are mean ± s.e.m. See Supplemental Table 4 for full statistical analysis.

**Figure 5.**
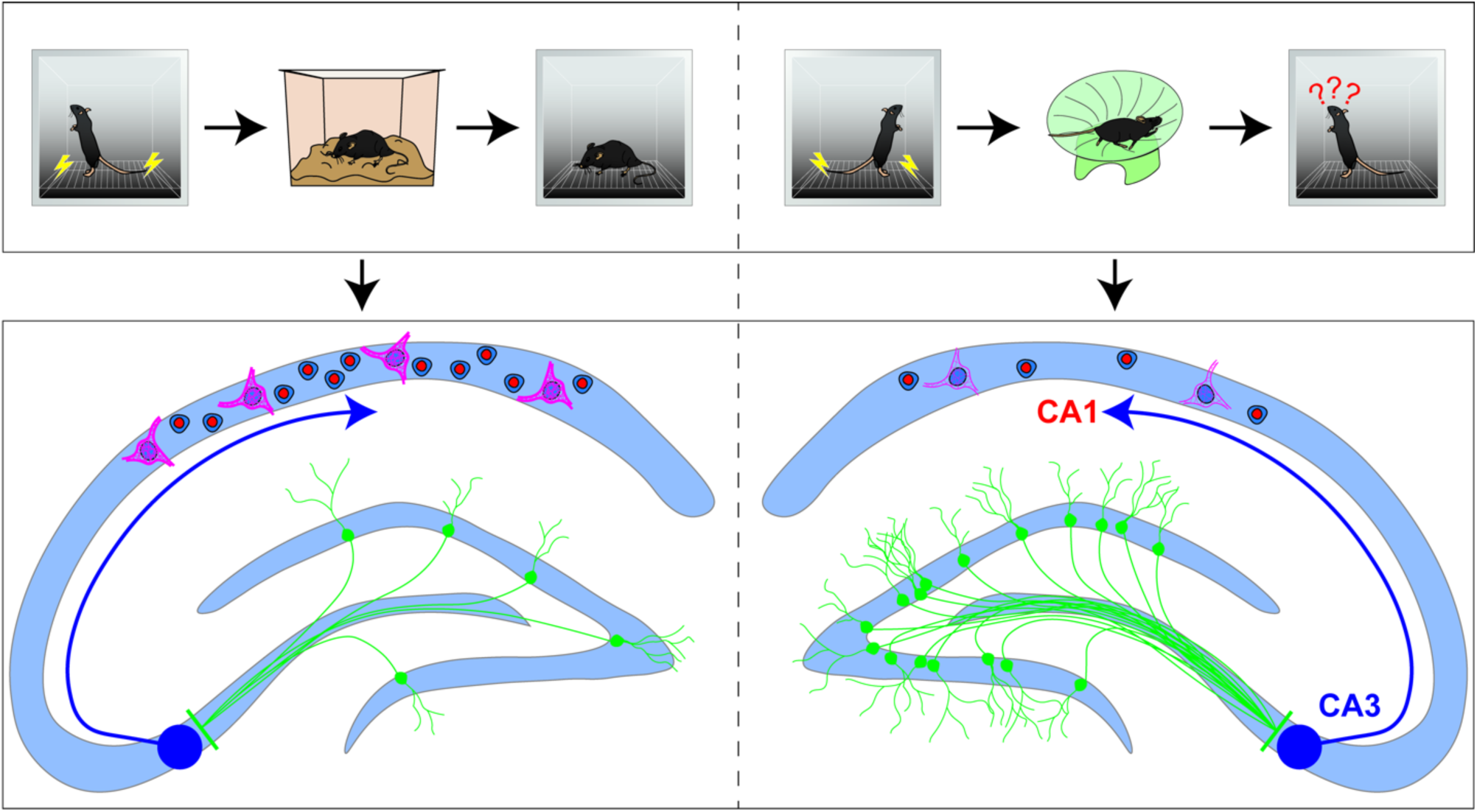
Summary of findings. Combined, these results show that increased neurogenesis causes large downstream effects in the activity and PNN expression of CA1 neurons and that, interestingly, the effects are largely absent in the DG and CA3 despite these areas being more proximal to adult-born granule cells. On this basis, we suggest that important mechanisms of neurogenesis-induced forgetting lie in physiological changes appearing distally and downstream in the CA1.

Adult neurogenesis in the hippocampus is strongly dependent on the age of the animal. Young mice have significantly greater levels of neurogenesis than do aged mice, and the decrease in neurogenesis begins early in life^13,38^. Approximately 1 month old is often considered to be the beginning of the period that constitutes adult neurogenesis^39^. The rate of neurogenesis drops precipitously beyond this time point, with significant decreases by as early as 2 months of age^40^. If the integration of adult generated neurons does in fact regulate CA1 PNN expression, then we would expect an increase in PNN expression with increasing age. To test this, we quantified PNN expression in CA1 of mice that were 1, 2 or 4 months old. As expected, we observed a significant age-dependent decrease in neurogenesis and a corresponding age-dependent increase in PNNs in CA1 (Fig. 4c,d)^41^. These results provide further supporting evidence that the levels of adult neurogenesis in the dentate gyrus impact CA1 PNN expression.

Together, the results of these various manipulations strongly suggest that adult neurogenesis modulated PNN expression in area CA1. But how does adult neurogenesis in the DG impact PNN expression in CA1ã Direct and/or local neuronal activity has previously been linked to modification of PNNs^42–44^. We hypothesized that the activity of adult-generated neurons could alter the circuit activity of the DG resulting in PNN changes in CA1. To examine the impact of new neuron activity, we used a Nestin-Cre mouse line to express ChR2 in immature neurons. Optogenetic stimulation of this immature neuron population was applied daily for 2 weeks beginning 24 hours after contextual fear conditioning. This optogenetic approach was sufficient to induce forgetting of the previously acquired contextual fear memory when tested one month after initial learning (Fig. 4e,f). The optogenetic stimulation significantly decreased PNN expression in CA1 compared to the non-stimulated control group (Fig. 4g). However, it did not increase the number of DCX+ cells in the DG (Fig. 4h). Together these findings demonstrate that the activity of immature neurons in the DG can have a direct impact on CA1 PNN expression.

## Discussion

We and others have demonstrated that elevated adult neurogenesis is inversely related to the strength of memories acquired before neurogenesis is increased^7–14^. Here, we sought to examine the mechanisms underlying this form of neurogenesis-induced forgetting. We began by identifying changes in experience-dependent c-Fos expression in the HPC associated with forgetting. We found that reduced contextual fear memory retrieval was accompanied by reduced c-Fos expression specifically within the CA1 subfield. This result points to the importance of CA1 in memory retrieval and suggests that adult neurogenesis disrupts memory storage and/or retrieval in this region. The CA1 has been shown to be active during recall of context fear memories^45,46^ and impairments in fear memory recall can be induced though optogenetic disruption of the CA1^47,48^. The CA1 has also been proposed by some to act not as a storage region *per se* but as a decoding circuit for memories stored in an orthogonalized fashion in CA3^49^. Regardless, of the exact function of the CA1, disruption of its activity clearly has critical implications for memory retrieval.

To further investigate the activity changes in CA1 associated with neurogenesis-induced forgetting, we performed *in vivo* fiber photometry. If increased neurogenesis (or alternatively non-specific effects of voluntary exercise) resulted in a generalized decrease in activity that blocked retrieval, then we would predict a decreased baseline Ca2+ signal in Ca1. However, we did not observe a baseline difference in activity but instead demonstrated that forgetting was associated with an overall context-specific reduction in CA1 population CA^2+^ signal as well as a behaviourally dependent decrease in peak frequency during movement. The decrease of these two proxies of neuronal activity in the CA1 during impaired memory retrieval suggests that the CA1 plays a critical role in the storage of context memories and that increased neurogenesis induces forgetting by modulating memory storage in CA1.

We next investigated neurogenesis induced changes in perineuronal nets in hippocampal subregions based on the proposed functions of these extracellular matrix components in modulating, plasticity and excitability^50^. We provide congruent evidence of the impact of positive modulators (running^8,9,11^, memantine^10^) and negative modulators (TMZ^8^, aging^40^) of adult neurogenesis on PNN expression in area CA1 of the hippocampus. While decreasing neurogenesis was associated with increased PNN expression within area CA1, increasing neurogenesis was associated with decreased PNN expression in CA1.

Prior evidence indicates that perineuronal nets in area CA1 protect the stability of long-term memories and, that their removal decreases memory storage through a mechanism involving increased activity of Parvalbumin expressing interneurons^34^. It has also been suggested that elimination of CA1 perineuronal nets acts to restrict long-term depression (LTD)^36,51^. Given that LTD can lead to the elimination of synapses^52^ this too could lead to a decrease in memory expression. Either, or both of these mechanisms could be induced by the reduction in CA1 perineuronal net expression that we identified from increasing the number of, or activity of adult generated neurons. These mechanisms are also consistent with decreased patterns of CA1 activity that we have observed here. If perineuronal net reduction increases the frequency of PV interneurons as has been previously shown, then a resulting decrease in the excitation of CA1 would be expected as was observed in terms of c-Fos expression and GCAMP7 activity.

Critically, we show here that direct optogenetic stimulation of immature adult generated neurons in the dentate gyrus both induced forgetting of a previously acquired memory and induces remodelling of the perineuronal nets in area CA1. This pattern replicates the behavioural phenomenon that we have observed with running as the modulator of adult neurogenesis. Previously it has been demonstrated using combined running and transgenic inhibition of adult neurogenesis, that the effects of running on forgetting of previously acquired memories are specifically dependent on the increase in neurogenesis. Our observation, that direct optogenetic stimulation of new neurons induces forgetting, provides further evidence of the specific effect of neurogenesis in this phenomenon.

Recent work has demonstrated that the density of perineuronal nets in the hippocampus can be bidirectionally modified by direct artificial stimulation or inhibition using DREADDs^44^. However, to the best of our knowledge our current results are among the first to demonstrate activity induced disruption of perineuronal nets occurring via a polysynaptic pathway rather than stimulation that is either localized to the region of interest or targeted to the PNNs themselves. This further emphasizes that although adult neurogenesis may modulate the structural connectivity of the local circuitry in the dentate gyrus, and proximally with CA3, the functional consequences of new neuron activity may be observed in areas that are not directly connected to the dentate gyrus. Previous research has found that new neurons in the DG preferentially incorporate with hilar interneurons and CA3 interneurons^22^. Thus, increased neurogenesis could have resulted in a shift toward more inhibitory drive from the DG onto the CA3, resulting in decreased c-Fos expression downstream of the DG. In addition, it has been shown that increased neurogenesis causes increased inhibition in the CA3 and CA1, with increased activity of parvalbumin interneurons in both regions and a decrease in CA1 sharp wave ripples^53^.

An explanation for why our c-fos and PNN effects appears largest in the CA1 may be that DG-CA3 connections are rather sparse compared with CA3-CA1 connections^23,24,26^, meaning in theory that a small change in population activity in the CA3 could have amplified downstream effects in the CA1.

Optimization of both learning and memory requires a balance between circuit plasticity and stability. Previous work has demonstrated that neurogenesis induced forgetting is accompanied by facilitation of new learning^9^. This multifaceted effect could be explained simply by an increase in plasticity which would shift the balance of the circuit to promote acquisition and disrupt memories that require the stability of that circuit for retrieval. Mechanistically, we provide strong evidence that adult neurogenesis alters this circuit balance through an activity dependent modulation of perineuronal nets in area CA1.

## Methods

### Subjects

We used two strains of mice. C57Bl/6N background mice (JAX) were used for most experiments. A transgenic mouse line expressing Cre under the nestin promoter sequence (JAX) was also used for optogenetic targeting of immature neurons. Female mice were used in all experiments and were 7 weeks old at the start of testing. Mice were housed in standard cages with three to five mice per cage and free access to food and water. The room lighting was maintained on a 12 hr /12 hr light/dark cycle (8 am, lights on). All behavioural testing and photometry experiments were conducted during the light cycle phase. Experiments were conducted in accordance with the policies and guidelines of the Canadian Council on Animal Care and were approved by the University of Calgary animal care committee.

### Manipulation of Neurogenesis

To increase neurogenesis, mice were given voluntary access to a running wheel (Fast Trac™, Med Associates Inc., Fairfax, VT, United States, 16 cm in diameter) in their home cage for one month. Voluntary running is a highly reproducible and widely used approach to increase adult neurogenesis. It was chosen here because it has been used in previous experiments examining neurogenesis-induced forgetting where the behavioural effects were found to be specifically dependent on the increase in neurogenesis ^8,9,11^. Mice were trained to use running wheels by gently placing them on the running wheel and obstructing their path off of the wheel. Mice were considered trained if they ran on the wheel for approximately 30 seconds. Wheel revolutions were recorded continuously using the Wheel Manager software program (Med Associates Inc., Fairfax, VT, United States) and the number of wheel rotations was divided by the number of mice in the cage to give an estimated running distance per mouse. Mice ran on average 8.1 km (± SEM 1.1 km) per day. Mice in the sedentary group were housed conventionally with no running wheel but had standard enrichment items (dome house and nesting material).

To quantify the effects of neurogenesis on perineuronal nets groups of mice were also treated with temozolomide (TMZ) to decrease neurogenesis or Memantine to increase neurogenesis. Control mice were injected with 0.9%Saline. TMZ was administered as previously described (REF). We injected mice on 3 consecutive days per week for 4 weeks with a dose of 25mg/kg (i.p.). Memantine was injected once per week for 4 weeks at a dose of 25 mg/kg (i.p.). Finally, we also used age as a naturalistic modulator of adult neurogenesis. C57bl/6N female mice were perfused at either 4, 8 or 12 weeks of age and perineuronal nets were labeled as described below.

### C-Fos Expression Experiment

#### Contextual fear conditioning

Prior to contextual fear conditioning, all mice were handled for five minutes each day for three to five days until mice were calm with the handler. The Ugo Basile (Gemonio, Italy) contextual fear conditioning chambers (17 cm x 17 cm x 24.7 cm) were placed in noise reducing cabinets. The conditioning environment had shock grid floors (bars spaced 0.5 cm apart, 0.2 cm in diameter). Behaviour was monitored using overhead infrared cameras and scored automatically using automated tracking software (ANY-Maze, Stoelting, Wood Dale, IL, United States). During the 5-minute training trial, mice were allowed to explore the chamber for two minutes before being delivered three shocks (1 mA, 2 s) each separated by an interval of 1 minute. Mice were removed from the conditioning chamber one minute after the last shock and placed back in their home cage undisturbed for 24 hours. Then, running wheels were introduced to the half of the mice. After four weeks of manipulating neurogenesis, mice were tested in the contextual fear task for five minutes with no shocks. The running wheel was locked the day prior to behavioural testing to prevent a direct effect of exercise on activity levels during testing. Freezing was used as the primary measure of memory retention and was defined as a complete lack of motion, except for respiration, for at least one second. Freezing was automatically calculated by the ANY-maze software system and was spot-checked for accuracy by an experimenter blind to treatment conditions. All mice were perfused 90 minutes following the context memory test in order to quantify c-Fos expression throughout the hippocampus.

### Pre-training Running experiment

In a separate experiment, mice were first given access to a running wheel for one month or housed without a wheel under standard conditions. The running wheels were then removed and the mice were trained on a contextual fear conditioning paradigm as described above. The mice were then tested one month later for context memory. All mice were perfused 90 minutes following the context memory test in order to quantify c-Fos expression throughout the hippocampus.

### Fiber Photometry Experiment

#### Surgeries

Surgeries were conducted under isoflurane anaesthesia delivered via a Somnosuite anaesthetic delivery system (Kent Scientific) drawing pure compressed oxygen. Mice were induced at 5% isoflurane concentration before being transferred to a stereotaxic head frame (Kopf) and maintained at ∼2% isoflurane. Internal body temperature was monitored with a rectal thermometer and regulated with a homeothermic pad. Analgesia (Anafen; 5 mg/kg; s.c) and fluid support (Lactated Ringer’s solution; s.c) were given at the beginning of the surgery. The scalp was shaved and cleaned with alternating chlorhexidine and 70% ethanol scrubs and then incised along the midline and the skull scrubbed with 3% H_2_O_2_. Using a robotic stereotaxic manipulator paired with Stereodrive software (Neurostar), the locations of bregma and lambda were measured as well as two points 2 mm to either side of the midline to allow the stereotaxic software to correct for any slight imperfections in skull flatness. After obtaining the coordinates of the injection and implant site, a burr hole was drilled in the skull (AP -2.18; ML ±2.1; DV to brain surface). A glass infusion needle attached to a Nanoject III infusion system (Drummond Scientific) and backfilled with mineral oil was slowly lowered into the CA1 (AP -2.18; ML ±2.0; DV 1.4) before injecting 250 nL of virus (pGP-AAV1-syn-jGCaMP7f-SV40-WPRE; Addgene) in 50 nL pulses each interspersed by ∼10 s and the final pulse followed by a 10 min period where the needle was left in place to allow for diffusion. At the end of the 10 min, the needle was gradually raised out of the brain before a fiber optic implant (N.A. 0.37, core 200µm 2 mm length; Neurophotometrics) was lowered into the CA1 just lateral to the injection site (AP -2.18; ML ±2.1; DV 1.4). Implants were secured to the skull by applying a thin layer of Metabond to the exposed surface of the skull and around the base of the implant. This layer was then covered with black opaque dental acrylic to form a headcap which was given time to harden before the incision was closed with suture material. Mice were then removed from the stereotaxic frame and allowed to recover for 2 weeks while receiving additional Anafen doses for 3 days after surgery.

#### Photometry recordings

Prior to behavioural training and *in vivo fiber photometry recordings*, mice were habituated to being connected to the fiber optic patch cord for 3 consecutive days. Fiber photometry was conducted using a FP3002 Neurophotometrics system controlled by Bonsai software^54^. The Neurophotometrics system also received TTL pulses generated by Anymaze software (Stoelting), which controlled the fear conditioning chambers, in order to synchronize Ca^2+^ recordings with the behaviour. Excitation light was delivered at 470 nm, while a 415 nm channel was used as an isosbestic control. Recordings were acquired at 40 FPS with the 2 channels being active in alternating frames, resulting in an effective framerate of 20/s for each channel. Light power was calibrated using a photometer (Thor Labs PM100D) so that each channel emitted at approximately 50 μW. During behaviour testing, the fiber patch cord (Doric) was secured to the mouse’s fiber implant with a ceramic collar and the mouse was left undisturbed with no recording for 30 s after the attachment of the patch cord. Following this, a 5 min baseline recording was acquired while the mouse sat alone in a clean transfer cage with bedding. At the 5 min mark, mice were transferred to the fear conditioning apparatus for training or testing.

#### Photometry Analysis

To synchronize photometry recordings with behavioural data, a TTL pulse was sent to the Neurophotometrics system upon the start of the ANY-maze protocol. Photometry recordings were analyzed using custom MATLAB scripts. Behavioural information recorded using ANY-maze was integrated into the analysis by indexing through the photometry data and aligning behavioural data based on the minimum difference between pairs of timestamps. Bouts of movement or immobility which were shorter than the freezing criteria of 2 seconds were not included in the analysis. Data from the isosbestic 410 nm channel was fit to a biexponential decay to correct for photobleaching ^55^. The resulting vector was used to linearly scale calcium-dependent data collected using a 470 nm LED ^55^. To calculate ΔF, these linearly scaled calcium-dependent data were subtracted from the raw unprocessed calcium-dependent data. The resulting values were divided by the linearly scaled calcium-dependent data to generate a ΔF/F trace.

For analyses of ΔF/F, traces were baseline corrected (Equation 1). This was done for each mouse by calculating the mean ΔF/F value during the middle 3 minutes of the 5-minute home cage recording. This period was chosen as the baseline period to reduce the likelihood that handling stress would influence the resulting value. The difference between the mean baseline ΔF/F and the minimum ΔF/F value recorded during the test session was then added to each value during the test. This correction was applied to account for traces in which the ΔF/F dropped below the X-axis during the test. The resulting ΔF/F traces were analyzed for area under the curve (AUC), peak frequency, and mean peak height. AUC was operationally defined as the summed area between the X-axis and the ΔF/F trace. Comparisons of AUC during specific behaviours were normalized by the summed duration of the specific behavioural epochs. Analyses of peak frequency and mean peak height employed a peak detection filter of 2 standard deviations above the median ΔF/F value of the session being analyzed ^56^.

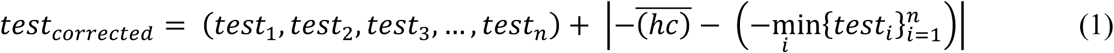

where ***hc*** is a vector representing the ΔF/F values during the home cage baseline sampling period, ***test*** is a vector representing the ΔF/F values during the testing period, and ***n*** represents the number of samples in the *test* vector. Equation (1) represents the calculation of the corrected ΔF/F during the test session.

### Optogenetic Manipulation Experiment

Nestin-cre mice underwent surgery using the surgical procedures similar to those described above. Here floxed channelrhodopsin expressing (AAV1-EF1a-double floxed-hChR2(H134R)-EYFP-WPRE-HGHpA) or GFP expressing (pAAV1-Ef1a-DIO EYFP) AAV viruses were infused bilaterally (0.25 µl) into the dentate gyrus (AP – 1.94, ML ± 1.25, DV 1.8). Fiber optic probes were implanted as described above. The mice were then left for 2 weeks to allow for recovery and viral expression. Mice were then injected with tamoxifen, 180 mg.kg per day) for 3 consecutive days and were then left for an additional 2 weeks before beginning behavioural experiments. Mice were first trained in the contextual fear conditioning paradigm as described above. Then, 24 hours following training mice received optogenetic stimulation with 0.4 mW, 10 HZ blue light for 1 minute, followed by 3 minutes of no stimulation. This was repeated consecutively 5 times for a total of 5 minutes of stimulation over a period of 20 minutes. This stimulation protocol was repeated daily for 2 weeks. One month following behavioural training and 2 weeks after the completion of optogenetic stimulation the mice were tested for contextual fear memory and were then perfused.

### Histology

#### Perfusions & Sectioning

Ninety minutes following the completion of the contextual conditioning task, mice were deeply anesthetized with isofluorane and then transcardially perfused with 0.1 M PBS and 4% formaldehyde. Brains were extracted and postfixed in 4% formaldehyde at 4°C for 24 hours. Brains were then cryoprotected with 30% sucrose in 0.1 M phosphate buffer at 4°C for three to five days until they sank. Brains were sectioned 40 μm thick on a cryostat (Leica CM 1950, Concord, ON, Canada) in 12 series for the c-Fos experiment and in 6 series for the fiber photometry experiment (in order to ensure that fiber tracks could be reliably located). Sections were kept in a buffered antifreeze solution containing 30% ethylene glycol, and 20% glycerol in 0.1M PBS and stored at -20°C.

#### Doublecortin labelling

Tissue sections were washed three times in 0.1 M PBS at room temperature for ten minutes each. Sections were then incubated for 48 h in a primary antibody solution containing either 1:500 goat anti-DCX antibody (c-18, sc-8066, Santa Cruz Biotechnology, Dallas, Texas, United States), 4% normal donkey serum, 0.4% Triton X, and 0.1 M PBS for the c-Fos experiment or, for the fiber photometry and PNN experiments, 1:200 rabbit anti-DCX antibody (Cell Signalling, 4604s), 3% normal goat serum, and 0.003% Triton-X in 0.1 M PBS. Sections then underwent three washes in 0.1 M PBS for ten minutes before being incubated for 24 h in a secondary antibody solution containing 1:500 Alexa Fluor 488 antibody (donkey anti-goat, CLAS10-1116, CedarLane Labs, Burlington, ON, Canada (or 1:500 Alexa Fluor 647 antibody goat anti-rabbit). Finally, sections were incubated with 1:5000 DAPI and 0.1 M PBS for 20 minutes and then washed twice in 0.1 M PBS for ten mins per wash. For each wash step, primary/secondary antibody solution, or DAPI incubation, the tissue was gently oscillated at room temperature. Sections were then mounted on glass slides and coverslipped with No. 1.5H coverslips using PVA-DABCO mounting medium.

#### Doublecortin Quantification

DCX labelled cells were counted in the granular and SGZ of the dentate gyrus of the hippocampus using an Olympus BX63 epifluorescent microscope and 60X oil immersion objective. Cells were counted if the cell was circular or ovular and the cell body could be clearly distinguished. Cells with a granular cytoplasmic pattern of fluorescence were excluded. DG area was quantified based on the DAPI counterstain by tracing the granule cell layer in each section using FIJI.

#### c-Fos labelling

Tissue sections were washed three times in 0.1 M PBS and then transferred to a primary antibody solution containing 1:500 rabbit anti-c-Fos antibody (226003, Synaptic Systems, Gottingen, Germany), 4% normal goat serum, 0.4% Triton X, and 0.1 M PBS for 48 h. Tissue sections were then washed in 0.1 M PBS three times for ten minutes before being transferred to a secondary antibody solution for 24 h. Secondary antibody solution contained 1:500 Alexa Fluor 488 antibody goat anti-rabbit (4412S, Cell Signaling Technologies, Danvers, MA, United States) with 0.1 M PBS. Finally, brain sections were incubated with 1:5000 DAPI and then washed twice in 0.1 M PBS. For each wash step, primary/secondary antibody solution, or DAPI incubation, the tissue was gently oscillated at room temperature. Sections were then mounted and coversliped with PVA-DABCO mounting medium.

#### c-Fos Image Acquisition

c-Fos labeled tissue was imaged using an Olympus FV3000 confocal microscope. All acquisition settings were kept constant across images. Images were acquired with a 10X objective and 2X optical zoom (total magnification of 20X), with the aperture set to 1 Airy unit. Z-stacks step size was 3.64 μm. Images were separated into DAPI and c-Fos channels and then maximum intensity projected as a 2D image using FIJI.

#### c-Fos Quantification

To minimize variability and bias in our c-Fos cell counts we validated and used a machine learning based approach (Supplemental Figure S1) Photomicrographs of c-Fos+ cells were segmented and converted to a binary image using the machine learning program Ilastik^57^ and quantified using FIJI. First, a training set of images which were representative of the experimental images were imported to Ilastik which was then trained to distinguish signal from background. This set of training images allowed Ilastik to classify cell from background based on pixel features including pixel intensity, edge features, and texture and object features such as object size, shape, and intensity. Following training, sample ROIs from 12 mice (five sedentary, seven running) from the DG, CA3 and CA1 region were uploaded to Ilastik to test the program. To compare the Ilastik-generated cell counts to ground truth values, the ROIs were also counted manually by four independent researchers who were experienced with quantifying c-Fos but were blind to the Ilastik-generated counts as well as to the counts of the other experimenters. The cell counts were summed for each region and then compared to Ilastik-generated counts. After training and validation, the experimental c-Fos images were batch processed. The binary images were imported to FIJI and overlaid with the DAPI image. Cells were then quantified using the binary image and area measured using FIJI. Area was then converted to mm^2^. Cells were quantified within the DG, CA3 and CA1 regions and expressed as a density (cells per mm^2^).

#### Wisteria Floribunda Lectin staining

PNNs were labelled via Wisteria Floribunda Lectin (WFA) staining. Cryoprotected sections were rinsed 3 times in 0.1M PBS and were then incubated in 0.2% triton-x in Carbo-free blocking buffer (VectorLABS) for 30 minutes. Then, the sections were stained in a 1:1000 dilution of FITC labeled WFA (VectorLABS) diluted in 1x carbo-free blocking buffer containing 0.05% tween-20 for 24 hours in the dark. Sections were then rinsed 3 times in PBS and counterstained with DAPI before being mounted and coverslipped with vectashield mounting medium (VectorLABS).

#### Quantification of Perineuronal Nets

WFA labelled perineuronal nets were manually counted in the granular zones of the dentate gyrus, area CA1, and area CA3 of the hippocampus. Counts were conducted using an Olympus BX63 epifluorescent microscope and 60X oil immersion objective by an experimenter blind to treatment condition. The areas of the granule zones of each region were quantified based on DAPI counterstaining. Granule cell layers were traced in each region from every section using CellSens (Olympus). Cells were quantified within the DG, CA3 and CA1 regions and expressed as a density (cells per mm^2^).

#### Quantification of CA1 Perineuronal Net Contiguity

Surface contiguity of CA1 perineuronal nets was assessed using an Olympus FV3000 confocal microscope. Image stacks of 5 randomly selected PNNs in the granular zone of area CA1 were collected from each mouse. Images were acquired with a 40X objective and 2.89X optical zoom (total magnification of 115.6X), with a numerical aperture of 0.95. Z-stacks step size was 0.37 μm. Imaging parameters were consistent across all scans. Image stacks were maximum intensity projected as 2D images using FIJI with the Z range set to encompass only the top half of each PNN. WFA signal was binarized in FIJI using default pixel intensity thresholding and individual PNNs were traced. Surface contiguity was defined as the fraction of WFA signal that resides within the largest connected component of the WFA signal after signal thresholding^58^.

#### Verification of Fiber Optic Placement and virus expression

Virus expression and fiber optic placement were verified by an experimenter blind to treatment condition and experimental outcomes. Virus expression in CA1 was present in all mice. Two control mice and three runners were removed from the photometry experiment because the tip of the optic fiber was positioned below CA1.

### Statistical Analysis

Statistical analysis was conducted using GraphPad Prism 8. Independent t-tests, two-way ANOVAs and Pearson Correlations were performed. Post-hoc Newman-Keuls post hoc tests were applied following ANOVA where appropriate. Grubbs’ analysis was utilized to identify any potential outliers in the Ilastik validation data. Hypothesis testing was complemented by estimation statistics for c-Fos quantification using estimationstats.com^59^. A multiple two-group analysis was used for c-Fos expression. For each two-group comparison, effect size (Cohen’s d) is calculated using a bootstrap sampling distribution using 5000 resamples along with a 95.0% confidence interval (CI; bias-corrected and accelerated). The data are plotted using Cumming estimation plots for multiple two-group analysis which show individual data points and the effect size of the comparisons. All statistical comparisons and outputs are included in supplemental tables 1-4.

## Acknowledgements

Funding for this study was provided by an NSERC Discovery Grant (RGPIN-2018-05135) to JRE and a Brain Canada Early Career research capacity building grant (4709) to JRE. DJT received a fellowship from the Canadian Open Neuroscience Platform. GAS received PDF fellowships from NSERC, the Hotchkiss Brain Institute and the Cumming School of Medicine.

## Author contributions

A.E., D.J.T., and J.R.E. conceived and designed the experiments. A.E., D.J.T., M.T. and G.A.S. performed the animal experiments.. A.E, D.J.T., G.A.S., and J.R.E. performed the histological procedures. A.E., D.J.T., and J.R.E. conducted the analyses. A.E., D.J.T., and J.R.E. wrote the paper.

## Data availability

The data that support the findings of this study are available in the manuscript or supplementary materials, and available from the corresponding author upon reasonable request.

## Code availability

All analysis code used to generate the results reported in the manuscript, instructions on how to use these analyses, and sample datasets have been made publicly available in a GitHub repository (https://github.com/dterstege/PublicationRepo/tree/main/Evans2021).

