## Supplemental Data for "Neurogenesis mediated plasticity is associated with reduced neuronal activity in CA1 during context fear memory retrieval"

**Supplemental Figure 1**

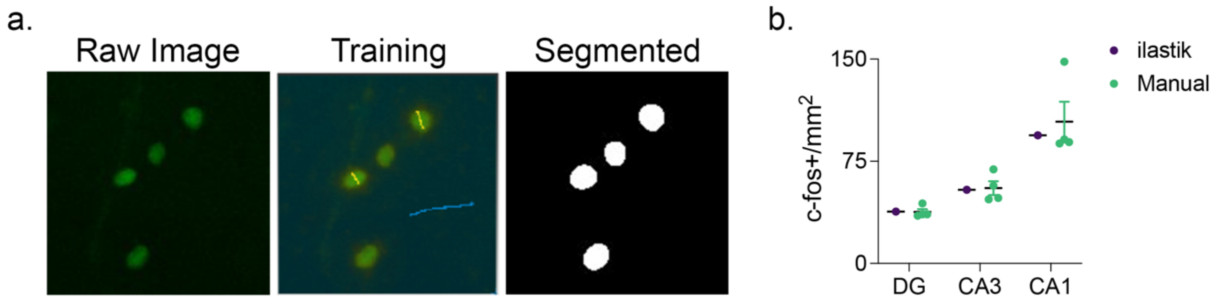

**Supplemental Figure 2**

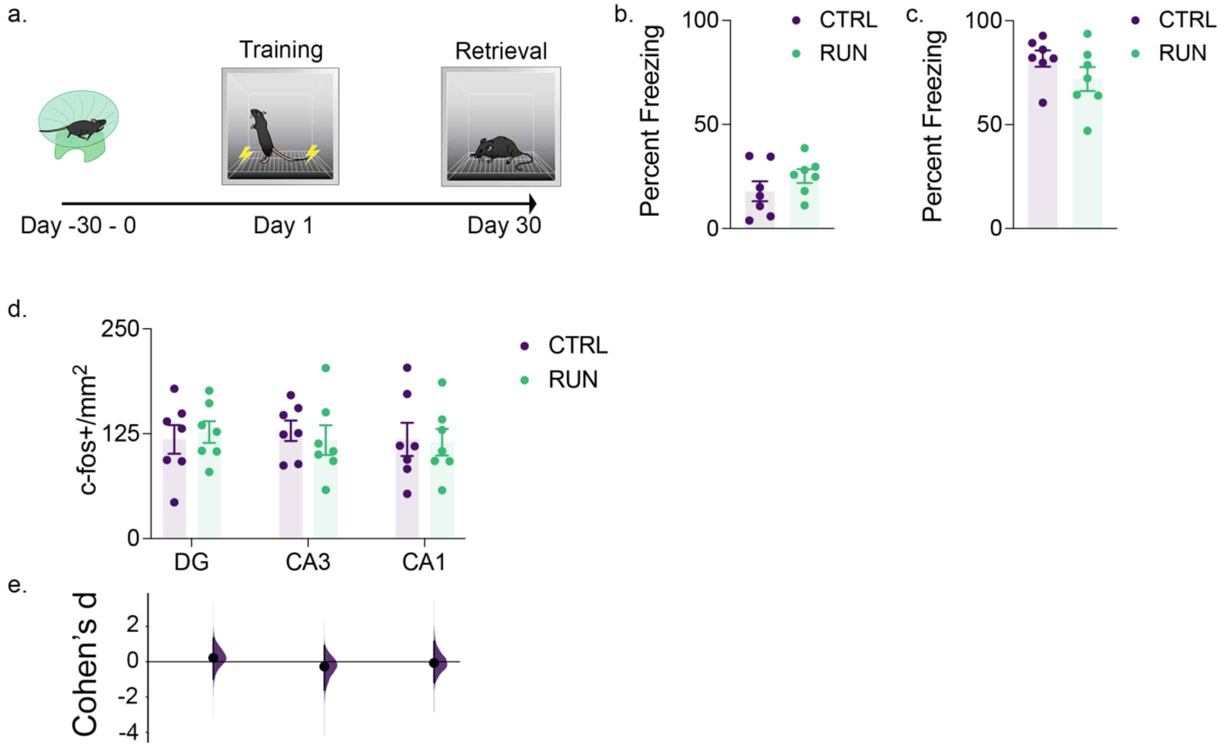

### Supplemental Figure 3

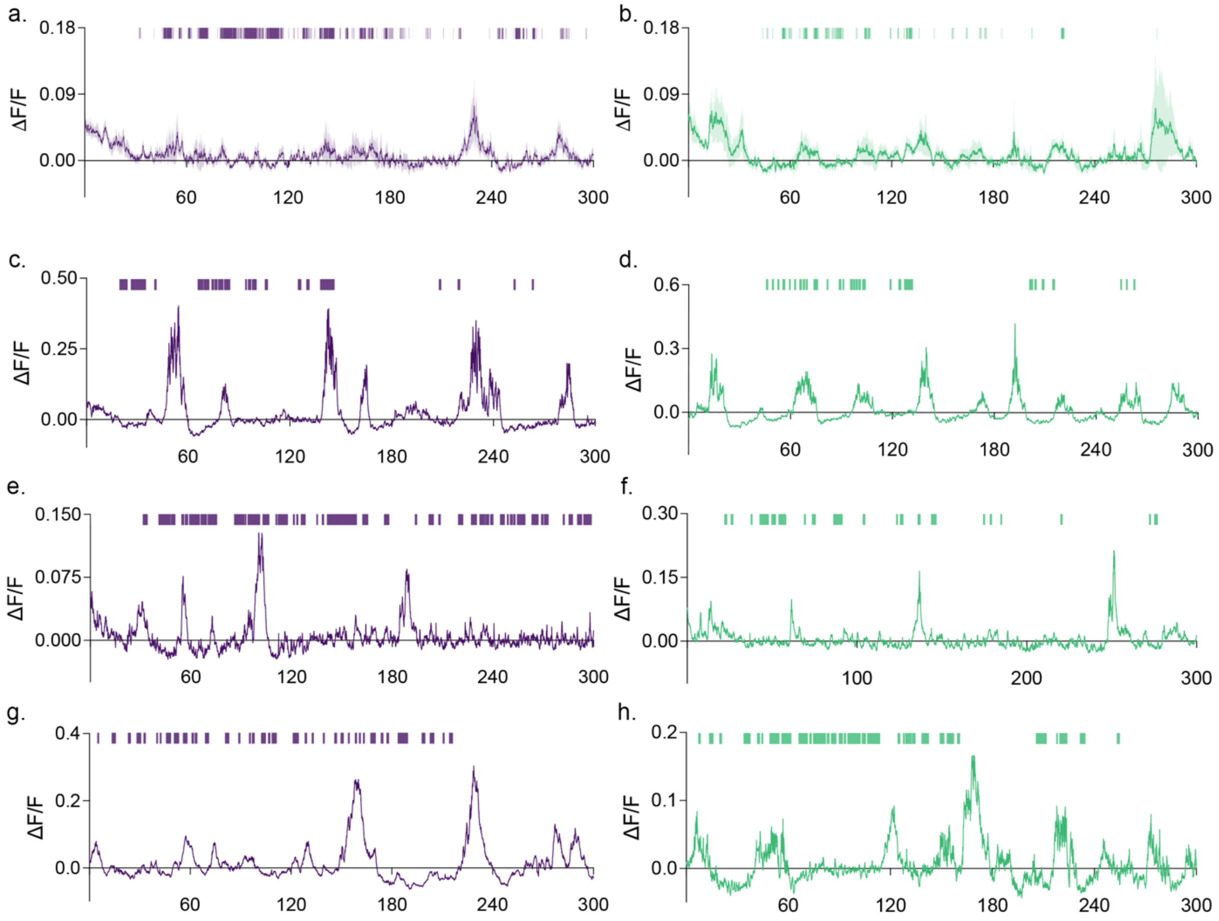

### Supplemental Figure 4

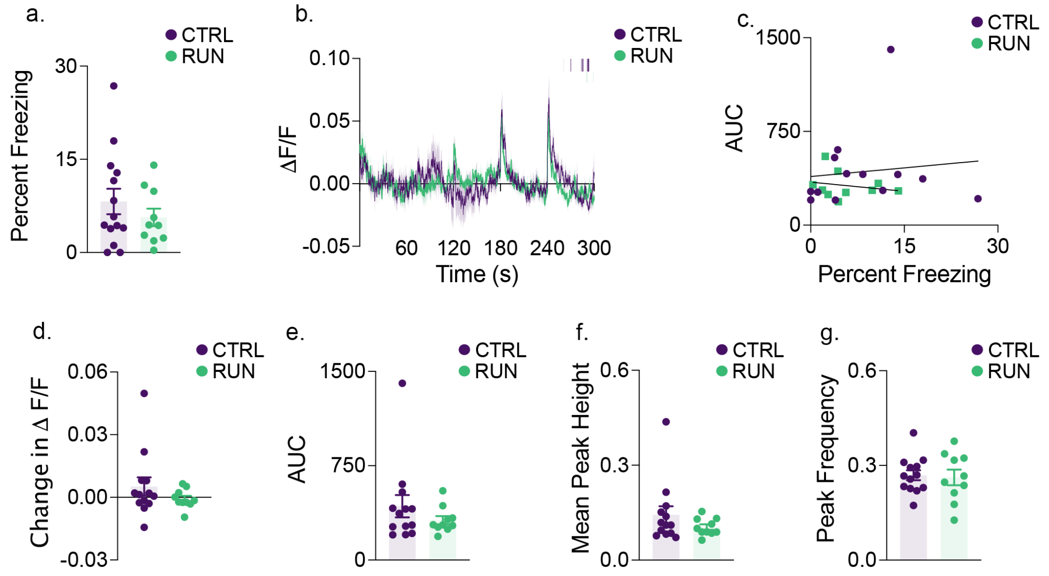

FIGURE CAPTIONS:

**Supplemental Figure S1 | Semi-automated c-fos+ cell segmentation using the supervised machine learning-based tool *ilastik.*** (**a**) Using supervised machine learning, the user interactively defined the criteria by which *ilastik* segmented c-fos+ cells from photomicrographs. (**b**) The segmentation of c-fos+ cells by *ilastik* yielded counts which were within the 95% confidence intervals of densities reported through independent hand scoring ($n$ = 4) of the same image sets in the dentate gyrus, CA3, and CA1. Data shown are mean ± 95% CI.

**Supplemental Figure S2 | The retrieval and underlying hippocampal activity of memories acquired after running-induced increased neurogenesis is not altered.** (**a**) After 30 days of running wheel access ($n$ = 7) or conventional housing ($n$ = 7), mice were contextually fear conditioned. Contextual memory retrieval was assessed 30 days later during context reintroduction. During both (**b**) training and (**c**) retrieval, prior running wheel access did not alter the proportion of the trial spent freezing. (**d**) The density of c-fos+ cells tagged to context reintroduction in the dentate gyrus, CA3, and CA1 did not change with prior running wheel access. (**e**) The magnitudes of effect sizes (Cohen's d) of the differences in c-fos+ cells in all three of the examined hippocampal subregions did not differ from a bootstrapped 95% confidence interval. Data analysis used Two-Sample T-Test (**b**,**c**), ANOVA (**d**) with Tukey's test during *post-hoc* multiple comparisons, and Multiple Two-Groups estimation statistics with Cohen's d as a measure of effect size (**h**). *$P < 0.05$. Data shown are mean ± s.e.m. See Supplemental Table 5 for full statistical analysis

**Supplemental Figure S3 | Non-baseline corrected group and representative individual recordings of CA1 activity during contextual memory retrieval.** Group mean traces from control (**a**, $n$ = 8) and runners (**b**, $n$ = 7) prior to baseline correction, with median group freezing plotted above. Representative individual traces and instantaneous freezing from control (**c**,**e**,**g**) and running (**d**,**f**,**h**) groups. Data shown are mean ± s.e.m.

**Supplemental Figure S4 | CA1 activity does not differ during contextual conditioning prior to running-induced changes in neurogenesis.** (**a**) During contextual fear conditioning prior to running wheel access ($n$ = 10) or conventional housing ($n$ = 14), there was no difference in the extent to which mice froze during the trial. (**b**) Mean GCaMP7f fluorescence across the context reintroduction trial with the mode of group freezing plotted above. (**c**) The area under the curve of the GCaMP7f fluorescence curve did not correlate with percent freezing in either group during this training trial. (**d**) When mice were transferred from the home cage to the conditioning chambers, neither group showed a significant change in mean GCaMP7f fluorescent signal. Throughout the training trial, neither group differed in (**e**) overall area under the GCaMP7f fluorescent curve, (**f**) mean photometry signal peak height, or (**g**) the frequency of peaks in the photometry signal. Data analysis used Two-Sample T-Test (**a**,**d,e,f,g**). *$P < 0.05$. Data shown are mean ± s.e.m. See Supplemental Table 6 for full statistical analysis

**Table 1: Statistics for the comparisons outlined in Figure 1.**

| **Two-Sample T Test, two-tailed** | | | | | |
|---|---|---|---|---|---|
| Figure | x-axis | y-axis | Groups (*n*) | p-value | t stat; df |
| **b** | Treatment Group | Percent Freezing | CTRL (14); RUN (14) | 0.1434 | t=1.509, df=26 |
| **c** | Treatment Group | Percent Freezing | CTRL (14); RUN (14) | 0.0005 | t=3.995, df=26 |
| **e** | Treatment Group | DCX+/mm2 | CTRL (14); RUN (14) | 0.0014 | t=3.574, df=26 |
| **Two-Factor ANOVA** | | | | | |
| Panel | x-axis | y-axis | Factor/*Comparison* | p-value | F stat; df |
| **g** | Region | c-fos+/mm2 | Interaction (*Tukey*) | 0.0114 | F (2, 72) = 4.768 |
| | | | *CA1:Sedentary vs. CA1:Running* | 0.0214 | |
| | | | *CA3 :Sedentary vs. CA3 :Running* | 0.6243 | |
| | | | *DG:Sedentary vs. DG:Running* | 0.884 | |
| | | | Areas of Hippocampus | <0.0001 | F (2, 72) = 12.09 |
| | | | Treatment | 0.0353 | F (1, 72) = 4.602 |

### Table 2: Statistics for the comparisons outlined in Figure 2.

| | | | | | |
|---|---|---|---|---|---|
| **Two-Sample T Test, two-tailed** | | | | | |
| Panel | x-axis | y-axis | Groups (*n*) | p-value | t stat; df |
| **c** | Treatment Group | DCX+/mm2 | CTRL (14); RUN (10) | 0.0417 | t=2.162, df=22 |
| **e** | Treatment Group | Percent Freezing | CTRL (14); RUN (10) | 0.0454 | t=2.121, df=22 |
| **g** | Treatment Group | ΔF/F | CTRL (14); RUN (10) | 0.9625 | t=0.04761, df=22 |
| **h** | Treatment Group | Change in ΔF/F | CTRL (14); RUN (10) | 0.0237 | t=2.429, df=22 |
| **Pearson Correlation** | | | | | |
| Panel | x-axis | y-axis | Groups (*n*) | p-value | correlation coefficient |
| **f** | Percent Freezing | DCX+/mm2 | CTRL (14) | 0.6457 | Pearson r=0.1349 |
| | | | RUN (10) | 0.0106 | Pearson r = -0.7610 |
| **j** | Percent Freezing | AUC | CTRL (14) | <0.0001 | Pearson r = 0.9980 |
| | | | RUN (10) | <0.0001 | Pearson r = 0.9933 |
| **Two-Factor ANOVA** | | | | | |
| Panel | x-axis | y-axis | Factor/*Comparison* | p-value | F stat; df |
| **k** | Behaviour | AUC | Interaction | 0.8959 | F (1, 44) = 0.01733 |
| | | | Behaviour | 0.9164 | F (1, 44) = 0.01116 |
| | | | Treatment Group | 0.0046 | F (1, 44) = 8.908 |
| **l** | Behaviour | Mean Peak Height | Interaction | 0.2856 | F (1, 44) = 1.168 |
| | | | Behaviour | 0.3867 | F (1, 44) = 0.7645 |
| | | | Treatment Group | 0.1135 | F (1, 44) = 2.607 |
| **m** | Behaviour | Peak Frequency | Interaction (*Tukey*) | 0.0039 | F (1, 44) = 9.297 |
| | | | *Freezing:CTRL vs. Moving:CTRL* | <0.0001 | |
| | | | *Freezing:RUN vs. Moving:RUN* | 0.0066 | |
| | | | *Moving:CTRL vs. Moving:RUN* | 0.0177 | |
| | | | *Freezing:CTRL vs. Freezing:RUN* | 0.6139 | |
| | | | Row Factor | 0.0009 | F (1, 44) = 12.72 |
| | | | Column Factor | 0.1952 | F (1, 44) = 1.730 |

### Table 3: Statistics for the comparisons outlined in Figure 3.

| **Two-Factor ANOVA** | | | | | |
|---|---|---|---|---|---|
| Panel | x-axis | y-axis | Factor/*Comparison* | p-value | F stat; df |
| **c** | Region | PNN+/mm2 | Interaction (*Tukey*) | 0.0842 | F (2, 39) = 2.638 |
| | | | *DG:CTRL vs. DG:RUN* | 0.9999 | |
| | | | *CA3:CTRL vs. CA3:RUN* | 0.9947 | |
| | | | *CA1:CTRL vs. CA1:RUN* | 0.031 | |
| | | | Region | <0.0001 | F (2, 39) = 59.95 |
| | | | Treatment Group | 0.0275 | F (1, 39) = 5.248 |
| **Two-Sample T Test, two-tailed** | | | | | |
| Panel | x-axis | y-axis | Groups (*n*) | p-value | t stat; df |
| **f** | Treatment Group | Contiguity | CTRL (8); RUN (7) | 0.0038 | t=3.511, df=13 |
| **Pearson Correlation** | | | | | |
| Panel | x-axis | y-axis | Groups (*n*) | p-value | correlation coefficient |
| **g** | DCX+/mm2 | CA1 PNN+/mm2 | CTRL (8) | 0.0844 | Pearson r=-0.6446 |
| | | | RUN (7) | 0.5586 | Pearson r=-0.2697 |
| **h** | DCX+/mm2 | Contiguity | CTRL (8) | 0.0423 | Pearson r=-0.7241 |
| | | | RUN (7) | 0.4492 | Pearson r=-0.3445 |

### Table 4: Statistics for the comparisons outlined in Figure 4.

| | | | | | |
|---|---|---|---|---|---|
| **Two-Factor ANOVA** | | | | | |
| Panel | x-axis | y-axis | Factor/*Comparison* | p-value | F stat; df |
| **a** | Region | PNN+/mm2 | Interaction (*Holm-Šídák*) | <0.0001 | F (4, 36) = 10.86 |
| | | | *CA1:TMZ vs. CA1:CTRL* | 0.0004 | |
| | | | *CA1:CTRL vs. CA1:MEM* | 0.0419 | |
| | | | *DG:TMZ vs. DG:CTRL* | 0.874 | |
| | | | *DG:CTRL vs. DG:MEM* | >0.9999 | |
| | | | *CA3:TMZ vs. CA3:CTRL* | 0.8216 | |
| | | | *CA3:CTRL vs. CA3:MEM* | 0.6997 | |
| | | | Region | <0.0001 | F (2, 36) = 21.87 |
| | | | Treatment Group | <0.0001 | F (2, 36) = 18.16 |
| **One-Factor ANOVA** | | | | | |
| Panel | x-axis | y-axis | Factor/*Comparison* | p-value | F stat; df |
| **b** | Treatment Group | DCX+/mm2 | ANOVA (*Holm-Šídák)* | 0.0009 | F (2, 12) = 13.39 |
| | | | *CTRL vs. TMZ* | 0.0366 | |
| | | | *CTRL vs MEM* | 0.0366 | |
| **c** | Treatment Group | PNN+/mm2 | ANOVA (*Tukey*) | 0.0005 | F (2, 11) = 16.78 |
| | | | *4 vs. 8* | 0.0161 | |
| | | | *4 vs. 12* | 0.0004 | |
| | | | *8 vs. 12* | 0.0198 | |
| **d** | Treatment Group | DCX+/mm2 | ANOVA (*Tukey*) | <0.0001 | F (2, 11) = 52.60 |
| | | | *4 vs. 8* | 0.0126 | |
| | | | *4 vs. 12* | <0.0001 | |
| | | | *8 vs. 12* | <0.0001 | |
| **Two-Sample T Test, two-tailed** | | | | | |
| Panel | x-axis | y-axis | Groups (*n*) | p-value | t stat; df |
| **f** | Treatment Group | Percent Freezing | CTRL (4); RUN (4) | 0.0190 | t=3.182, df=6 |
| **g** | Treatment Group | PNN+/mm2 | CTRL (4); RUN (4) | <0.0001 | t=9.937, df=6 |
| **h** | Treatment Group | DCX+/mm2 | CTRL (4); RUN (4) | 0.9259 | t=0.09694, df=6 |

### Table 5: Statistics for the comparisons outlined in Supplemental Figure S2.

| **Two-Sample T Test, two-tailed** | | | | | |
|---|---|---|---|---|---|
| Panel | x-axis | y-axis | Groups (*n*) | p-value | t stat; df |
| **b** | Treatment Group | Percent Freezing | CTRL (7); RUN (7) | 0.2344 | t=1.252, df=12 |
| **c** | Treatment Group | Percent Freezing | CTRL (7); RUN (7) | 0.185 | t=1.406, df=12 |
| **Two-Factor ANOVA** | | | | | |
| Panel | x-axis | y-axis | Factor/Comparison | p-value | F stat; df |
| **d** | Region | c-fos+/mm2 | Interaction | 0.8394 | F (3, 48) = 0.2802 |
| | | | Region | 0.908 | F (3, 48) = 0.1822 |
| | | | Treatment Group | 0.7853 | F (1, 48) = 0.07507 |

### Table 6: Statistics for the comparisons outlined in Supplemental Figure S3.

| **Two-Sample T Test, two-tailed** | | | | | |
|---|---|---|---|---|---|
| Panel | x-axis | y-axis | Groups (*n*) | p-value | t stat; df |
| **a** | Treatment Group | Percent Freezing | CTRL (14); RUN (10) | 0.3574 | t=0.9401, df=22 |
| **d** | Treatment Group | Change in ΔF/F | CTRL (13); RUN (10) | 0.2685 | t=1.137, df=21 |
| **e** | Treatment Group | AUC | CTRL (13); RUN (10) | 0.3031 | t=1.056, df=21 |
| **f** | Treatment Group | Mean Peak Height | CTRL (13); RUN (10) | 0.2358 | t=1.221, df=21 |
| **g** | Treatment Group | Peak Frequency | CTRL (13); RUN (10) | 0.8067 | t=0.2478, df=21 |
| **Pearson Correlation** | | | | | |
| Panel | x-axis | y-axis | Groups (*n*) | p-value | correlation coefficient |
| **c** | Percent Freezing | AUC | CTRL (13) | 0.7107 | Pearson r=-0.1140 |
| | Percent Freezing | AUC | RUN (10) | 0.5615 | Pearson r=-0.2094 |
